# Inferring critical points of ecosystem transitions from spatial data

**DOI:** 10.1101/187799

**Authors:** Sabiha Majumder, Krishnapriya Tamma, Sriram Ramaswamy, Vishwesha Guttal

**Affiliations:** Department of Physics, Indian Institute of Science, Bengaluru 560 012, India; Centre for Ecological Sciences, Indian Institute of Science, Bengaluru 560 012, India

## Abstract

Ecosystems can undergo abrupt transitions from one state to an alternative stable state when the driver crosses a threshold or a critical point. Dynamical systems theory suggests that systems take long to recover from perturbations near such transitions. This leads to characteristic changes in the dynamics of the system, which can be used as early warning signals of imminent transitions. However, these signals are qualitative and cannot quantify the critical points. Here, we propose a method to estimate critical points quantitatively from spatial data. We employ a spatial model of vegetation that shows a transition from vegetated to bare state. We show that the critical point can be estimated as the ecosystem state and the driver values at which spatial variance and autocorrelation are maximum. We demonstrate the validity of this method by analysing spatial data from regions of Africa and Australia that exhibit alternative vegetation biomes.

## 1 Introduction

Many ecosystems, ranging from tropical forests to coral reefs, are stable across a range of environmental conditions. However, when a control parameter or driver in the environment crosses a threshold value, ecosystems may undergo abrupt shifts from their current state to an alternative state (Noy-Meir 1975; Scheffer *et al.* 2001; Hughes 1994; van de Koppel *et al.* 1997; Steele 1998). This threshold is called a critical point or, in the dynamical systems literature, a bifurcation. These drastic changes of state, termed critical transitions, may result in loss of biodiversity and ecosystem services. Therefore, estimating critical points and locating ecosystems’ current parameters relative to such critical values are matters of importance.

Estimating critical points of real ecosystems can be notoriously difficult. One novel approach relies on the analysis of dynamics of state variables following large perturbations (D’Souza *et al.* 2015); however, experimentally induced large perturbations can push the system to a transition. Critical points can also be estimated from long-term data of ecosystems that have previously undergone transitions (Ratajczak *et al.* 2014). Alternatively, one could construct a complete characterization of ecosystem states as a function of drivers (Hirota *et al.* 2011; Staver *et al.* 2011; Staal *et al.* 2016), and estimate critical points. However, these methods are limited to a few ecosystems where data are available in steady-state conditions and over large enough spatial or temporal scales, thus limiting their applicability. Therefore, most recent studies have focused on obtaining qualitative ‘early warning signals’ (EWS) of critical transitions which are based on theories of dynamical systems and phase transitions (Wissel 1984; Scheffer *et al.* 2009; Carpenter & Brock 2006; Guttal & Jayaprakash 2008; Van Nes & Scheffer 2007).

These EWS have been empirically tested in aquatic, savanna and climatic systems (Carpenter *et al.* 2011; Eby *et al.* 2017; Dakos *et al.* 2008). Various studies have also highlighted their limitations (Boettiger & Hastings 2013), for example, due to insufficient sampling, stochasticity or short length of ecological datasets (Perretti & Munch 2012; Guttal *et al.* 2016; Burthe *et al.* 2016). Even reliable measurements of these signals do not provide quantitative estimates of how far the system parameters are from the critical point.

The goal of our manuscript is to develop a method that offers quantitative estimates of critical points. We hypothesise that values of drivers and state variables in regions with maximum spatial variance and autocorrelation of ecosystem states offer estimates of critical point (Box 1). We argue that this method is applicable even if data are not in steady-state conditions and are available only at relatively small scales than those required for a complete characterisation of ecosystem states. We justify our claim based on analyses of models and remotely-sensed vegetation data from Africa and Australia. Therefore, one can employ transects that span alternative stable states of ecosystems. The parameters of biomass density, grazing or rainfall values at which maxima of spatial metrics occur offer approximations of the critical points.

## 2 Box 1: Maxima of spatial variance and autocorrelation occur at the critical point

To see why spatial variance and autocorrelation are maximum at the critical point, we consider a generic model of abrupt transition given by

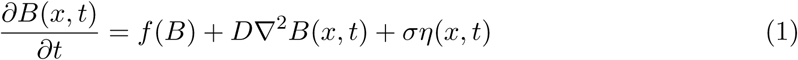

where *f*(*B*) represents local growth rate of population density *B*, the second term (diffusion) represents spatial interactions and the third term represents Gaussian fluctuations, uncorrelated in space (*x*) and time (*t*), with a strength *σ*. We assume *f*(*B*) such that the nonspatial and deterministic version of Eq (1) exhibits multiple stable states and a saddle-node bifurcation. We investigate the dynamics for the spatially-extended case in a simplified analytical approach, working in one space dimension and linearising the system in the vicinity of a stable state, to obtain

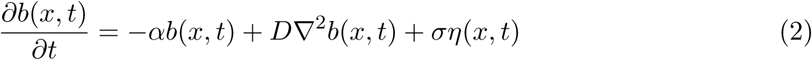

where 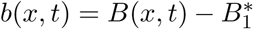 and *α* measures the distance from the critical point. The spatial variance 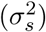 and autocorrelation (*ACF*) of this system variable (*b*) are easily found to be:

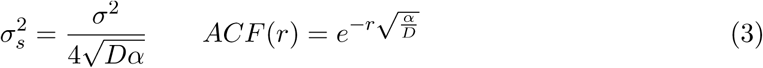

Thus, approaching the critical point (*α* → 0) gives rise to enhanced fluctuations of the ecosystem state and extended correlations in space (Guttal 2008; Guttal & Jayaprakash 2009; Dakos *et al.* 2010). At the critical point (*α* = 0), within our approximation, 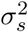 is infinite and ACF(*r*) becomes unity for all *r* (Scheffer *et al.* 2009). These extreme results arise can be traced to the linearised approximation which is well known (Chaikin & Lubensky (2000), chapter 5) to exaggerate fluctuation magnitudes for one‐ or two-dimensional systems. An exact calculation based on the theory of critical phenomena (Chaikin & Lubensky 2000) shows that 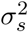 remains finite, ACF decays with *r*, but both attain a maximum at the critical point.

Based on these observations, we hypothesise that maxima in both spatial variance and spatial autocorrelation of the state indicates that the system is at the critical point of a transition.

## 3 Material and methods

### 3.1 Analyses of spatially-explicit models to identify critical points

#### Spatially-explicit model showing critical transitions

In our model, the landscape contains *N* × *N* cells, with each cell in a state of being empty (0) or occupied by a plant (1). In the simplest version of this model, known as contact process (Durrett & Neuhauser 1991), a focal plant germinates a nearby empty cell with probability *p*, or dies with a probability 1 − *p*. This model exhibits a transition from a bare state (*p* < *p*_*c*_) to vegetated state (*p* ≥ *p*_*c*_) with the critical point *p*_*c*_ ≈ 0.62. This transition is, however, *continuous*. We therefore consider an extended version of the model (Lübeck 2006). Here, the baseline birth (*p*) and death (1 − *p*) probabilities are modified via a local positive feedback (denoted by *q*), as an increase in birth probability and a decrease in death probability for plants which are surrounded by other plants. We assume that parameters *p* and *q* do not vary across cells and hence call this a ‘homogeneous-driver model’. This model with large *q* exhibits a *discontinuous* transition from a vegetated state (*ρ*_*c*_ = 0.32) to bare state at a critical value of the driver (*p*_*c*_ = 0.2852; see Fig. S2 in Appendix S2 in Supporting information).

#### Model with a gradient of driver along space

To reflect real-world situations where drivers such as rainfall, grazing or fire are spatially heterogeneous, we consider a simple case where the driver changes from low to high values along one dimension of a two dimensional landscape; for instance, this may represent rainfall gradients observed in tropical forest biomes or savanna ecosystems (Favier *et al.* 2012; Eby *et al.* 2017). Hence, we model the landscape as a rectangular matrix of width *N* and length *N* × *l* with a homogeneous positive feedback (*q*). However, the baseline birth probability (*p*) increases from *p*_*l*_ to *p*_*h*_ along the length of the matrix such that the system in its steady state exhibits a transition in this range of *p* (Fig. S4 in Appendix S2). We call this a ‘gradient-driver model’. Since real-world ecosystems are rarely in steady state, we stopped the simulation at 1500 time steps; this is in contrast to steady-state simulations that require 1 million time steps near critical points.

#### Null model

We use a null model from (Kéfi *et al.* 2011) where both birth (*p*) and death (*d*) probabilities are independent of the state of the neighbouring cells. Here, vegetation density reduces gradually as a function of reducing *p*, with no critical points.

#### Computing spatial metrics

We compute spatial variance and spatial autocorrelation at lag-1 using methods from Kéfi *et al.* (2014); Sankaran *et al.* (2017). Studies show that spatial variance in binary-state spatial data (e.g. occupied (1) or empty (0) at each location) depends only on mean cover and does not capture spatial structure of the data (Eby *et al.* 2017; Sankaran *et al.* 2017). One must average spatial data over local spatial scales, known as ‘coarse-graining’ (Sethna 2006), before computing spatial metrics. To restate our hypothesis in the context of this method, we expect spatial variance (referred to as variance method) and spatial ACF-1 (autocorrelation at lag 1, referred to as ACF method) to be maximum at critical points if the data are optimally coarse-grained. See Appendix S1 for formula for spatial metrics, details on coarse-graining spatial data by a scale *l*_*cg*_ and a method to obtain an ‘optimal coarse-graining length 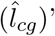 at which critical points can be estimated.

### 3.2 Validation of the method using real data

To demonstrate the empirical validity of our method, we used vegetation data from three regions as shown in Fig. 2. We first estimate critical points from our method at the relatively small spatial scale of transects (8 km *×* 90 km) that span alternative stable states of vegetation. We then compare these estimated values to those from an independent method at a larger *landscape scale* from these regions (∼ 200 km × 250 km).

**Figure 1:**
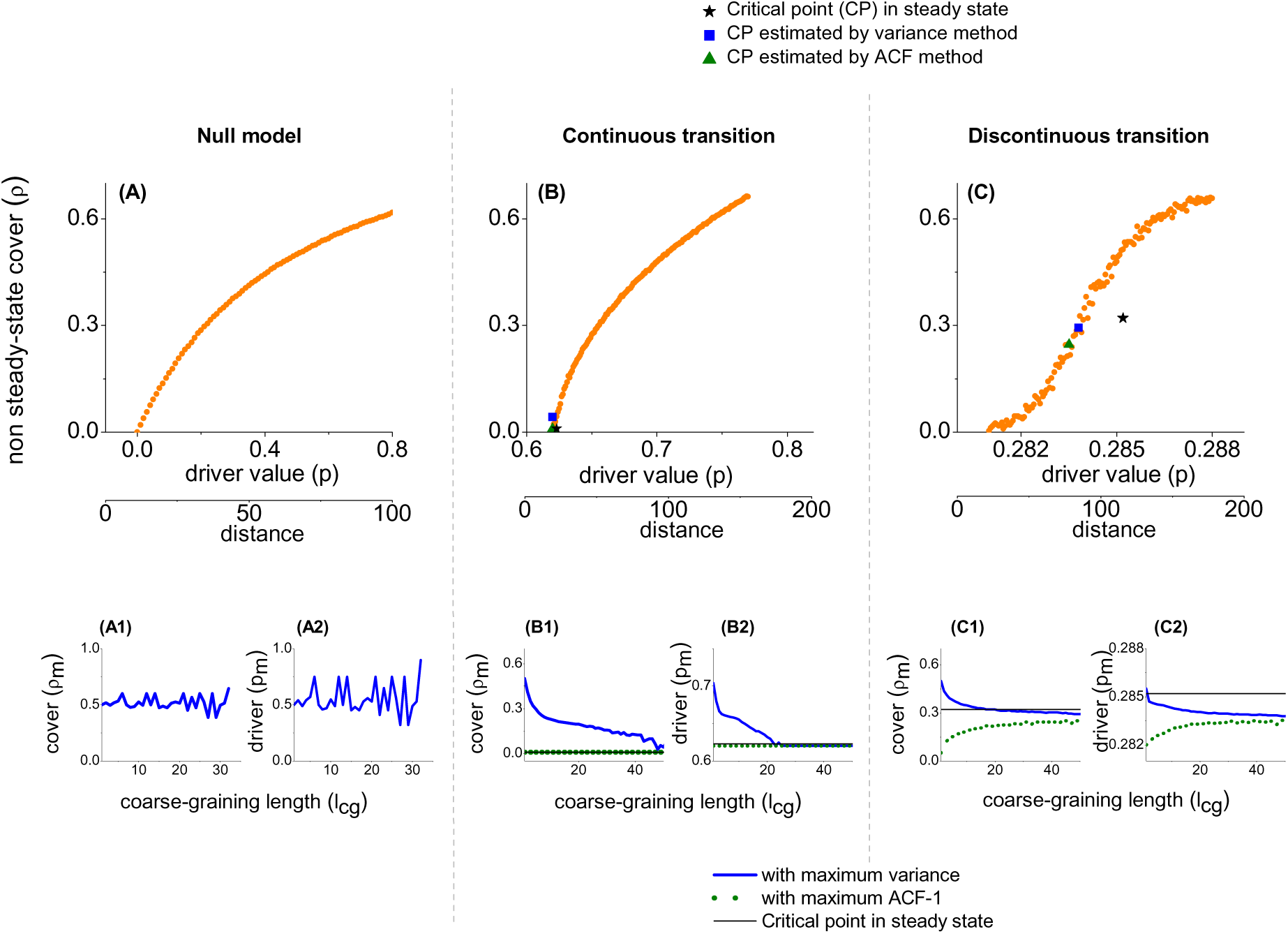
Simulations of spatially-explicit ecological models show that even for data arising from nonsteady state and gradient-driver conditions, estimated critical points (blue squares and green triangles in B and C) are reasonably close to the critical point in steady states (black star). In the null model with no critical points (A), as theoretically expected, the peak of spatial variance occurs around density of 0.5 for all coarse-graining lengths (A1, A2), but there is no peak for spatial ACF-1 (see Fig. S2, S3 in Appendix S2). Thus, we infer there is no critical point for the null model. In the continuous transition model (B), peaks of spatial variance and ACF-1 (B1, B2) converge close to the steady state critical points. In (C, C1 and C2), we find qualitatively similar results for the discontinuous transition model.

**Figure 2:**
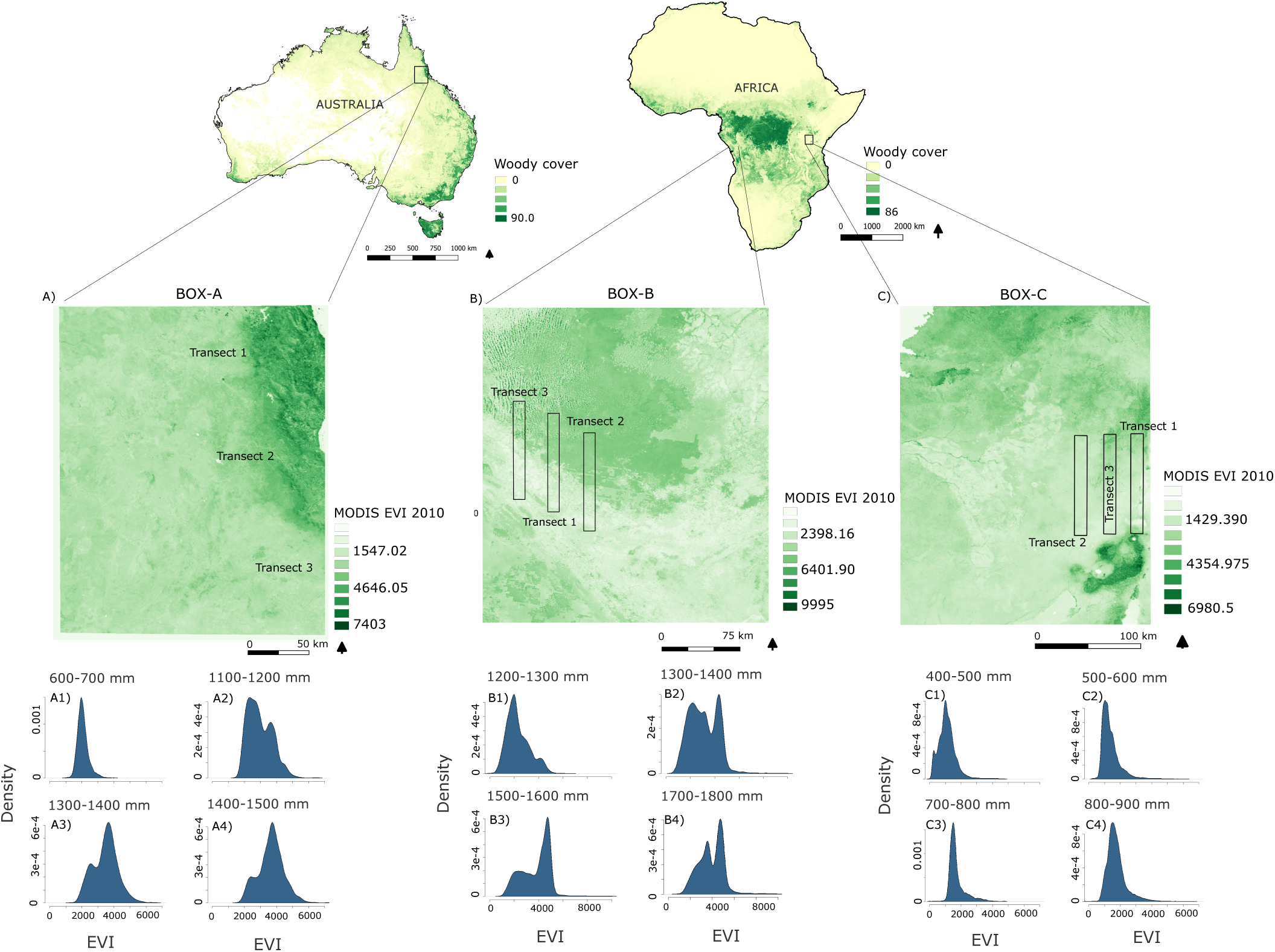
Location of study sites and the transects. (A, B and C) show the spatial distribution of EVI in Box-A (Australia), Box-B (Congo-Gabon in Africa) and Box-C (Serengeti in Africa) respectively. (A1-A4 and B1-B4) show that EVI changes from a unimodal to a bimodal frequency distribution as a function of mean annual rainfall within Box A and B. (C1-C4) show that EVI distributions remain unimodal for all the rainfall ranges in Box-C.

#### Study sites

We use results from Staver *et al.* (2011) to find regions (∼200 km *×* 250 km) that show bistable states of forests and grasslands (Appendix S3). We choose two regions, one in Australia (Box-A shown in Fig. 2) and one at the Congo-Gabon border in Africa (Box-B), where the vegetation cover varies from high (∼ 70%) to low (∼ 20%). We also select a region including Serengeti National Park (Box-C), which shows low vegetation cover (< 35%).

#### Vegetation and Rainfall Data

Remotely-sensed vegetation indices, such as Normalized Difference Vegetation Index (NDVI) and Enhanced Vegetation Index (EVI), are related to the amount of photosynthetic activity. We employ EVI as a proxy for vegetation cover since, unlike NDVI, it does not saturate at high values of photosynthetic activity and thus is a better proxy for both low and high vegetation covers (Glenn *et al.* 2008). We obtain EVI data from the Moderate Resolution Imaging Spectroradiometer (MODIS; using Google Earth Engine platform (Google Earth Engine Team 2015) for 2010 at 250 m resolution (Huete *et al.* 2002). We choose the dry months (June - August) to minimise cloud cover. We analyse the relation between rainfall (WorldClim, 1 km resolution) (Hijmans *et al.* 2005)) and vegetation cover as rainfall is an important driver of vegetation (Sankaran *et al.* 2008; Hirota *et al.* 2011; Staver *et al.* 2011).

#### Estimating critical points from transects

We construct 8 km *×* 90 km transects, which are ∼ 1.5% of area of Boxes, within each landscape such that they capture the gradient in EVI (Fig. 2). Transect-3 in Box-A is limited to 70km to avoid regions of high human activity, and Transect-1 in Box-C to 60km to avoid steep changes in altitude.

For each transect, we employ a moving window of size 8 km *×* 8 km, with a moving distance of 2km along its length. We calculate spatial variance and ACF-1 of coarse-grained EVI data along the moving window. We smooth the spatial-metrics data using ‘smooth.spline’ in R (v 3.3.1) and identify peaks as local maxima of the smoothed function. If the peaks in spatial variance and spatial ACF-1 occur within a distance of 4 km, we define them as coinciding peaks. To test if the coincidence of the peaks can occur by chance, we compute the spatial metrics on null transect data, which is obtained by shuffling EVI data for each moving window of the transect. We also test if the peaks are likely to be caused by landscape heterogeneity (see below). We hypothesise that coinciding peaks in spatial variance and ACF-1 in the absence of such confounding effects correspond to the critical points. See Appendix S5 for a detailed step-by-step procedure.

#### Confounding factors: Human influence and landscape heterogeneity

We only choose regions with minimal human influence (using the 2009 GlobCover maps (Bontemps *et al.* 2011)). The observed patterns of spatial metrics in vegetation transects may also arise from landscape heterogeneity, such as elevation, aspect, slope and soil variations (Reed *et al.* 2009), rather than as a consequence of the underlying dynamics. We compute landscape heterogeneity (calculated as spatial variance in elevation, slope, aspect, and soil-type richness) along each transect and identify its peaks, referred as heterogeneity peaks, based on the smoothing procedure described previously. We then reject peaks in spatial variance and ACF-1 if they occur within 4 km of any of the heterogeneity peaks.

#### Multimodality in vegetation cover and estimation of critical points from landscape analyses

We divide rainfall into 100 mm bins (Hirota *et al.* 2011). For each bin, we smooth the frequency distribution of EVI using the function ‘density’ in R (v 3.3.1) and identify modes as local maxima of the density functions (Appendix S4). We test whether the observed multimodality in EVI is associated with multimodality in rainfall, and reject such modes because they are unlikely to reflect alternative stable states in EVI. Now, from the location of remaining modes, we obtain an independent estimate of critical points, defined as the rainfall levels at which EVI distributions change from bimodal to unimodal. We refer to the resulting plots of modes vs rainfall as ‘state diagrams’.

## 4 Results

### 4.1 Estimating critical points in models

#### Spatial variance and autocorrelation can estimate critical points in the models

We apply our method to three scenarios of the gradient-driver model (Fig. 1). In steady-state conditions, they correspond to qualitatively different behaviours, i.e. no transition, a continuous transition and a discontinuous transition, respectively (Fig. S4 in Appendix S2). In nonsteady-state conditions, however, the analysis of driver-state relationships alone (Fig. 1 A, B or C), may not be sufficient to yield critical points. This is because vegetation cover changes gradually from large values to the bare state in all three cases. As we describe below, our methods provide estimates that are reasonably close to the steady-state critical points even when applied to nonsteady-state data.

Spatial variance for raw data (i.e. without coarse-graining, *l*_*cg*_ = 1) shows a maximum value for the snapshot with 50% cover (Fig. 1 A1, B1, C1), which is not the critical value of cover in our models. This is expected, as argued previously (Eby *et al.* 2017; Sankaran *et al.* 2017), in binary-valued spatial data. In the null model, the peak of variance does not change with *l_cg_* (Fig. 1 A1, A2). In contrast, for both continuous and discontinuous transition models, after coarse-graining, the values of cover and driver with maximum spatial variance, denoted as *ρ_m_* and *p_m_* respectively, change with increasing *l_cg_*, converging to the steady-state critical-point values *ρ_c_* and *p*_*c*_ (Fig. 1 B1, B2, C1, C2). Likewise, patterns of spatial ACF-1 differ between the null model and the other two models with transitions, with the null model showing no peak as a function of density.

Due to the lack of match in the patterns of peaks of spatial variance and ACF-1, we conclude that the null model, as expected, has no critical points. In contrast, for the continuous transition scenario, the variance and the ACF methods yield critical points of (*ρ*, *p*)=(0, 0.63) and (0, 0.623) respectively. These values are close to the steady-state critical point (*ρ*_*c*_, *p*_*c*_)=(0, 0.623). Likewise, for the discontinuous transition scenario, the variance and the ACF methods yield estimates of (*ρ*, *p*)=(0.30, 0.2838) and (0.28, 0.2835) which are reassuringly close to the actual critical point in steady state (*ρ*_*c*_,*p*_*c*_)= (0.32, 0.2851).

Our method can also estimate critical points in models of semi-arid vegetation that incorporate complex and detailed ecological processes (Kéfi *et al.* 2007; Schneider & Kéfi 2015) (Fig. S5 in Appendix S2). These findings provide a proof of principle, encouraging us to apply this method to real world data.

### 4.2 Application to find critical points in real ecosystems

#### Estimation of critical points from transects

We show the results of one representative transect from each box in Fig. 3 and others are shown in Fig. S13, S14, S15 in Appendix S5. For all transects, EVI typically increases with rainfall (Fig. 3 A1-C1). We find multiple peaks of spatial variance and ACF-1 in EVI for these transects. We reject peaks that occur in the vicinity of the landscape heterogeneity peaks (grey bands). Rainfall values corresponding to coinciding peaks of spatial variance and ACF-1, 1108 - 1334 mm/year for Box-A and 1281 - 1306 mm/year for Box-B (Table 1), are estimated as critical points of the transition. For Box-C, after accounting for heterogeneity, we found no coinciding peaks, and hence no critical point estimations, in any of the transects.

**Figure 3:**
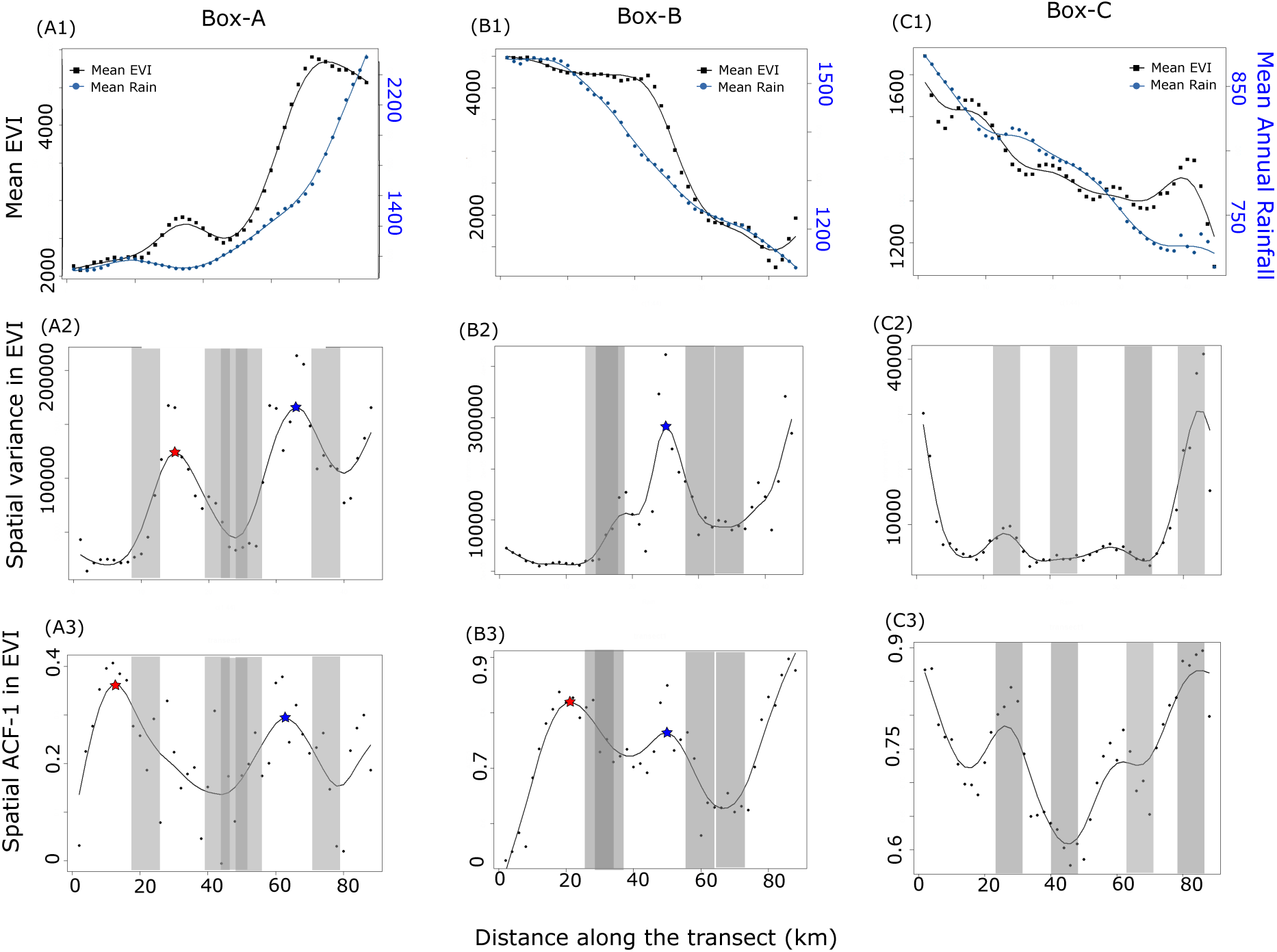
Estimation of critical points (blue stars in A2, B2, A3, B3) from the analyses of transects after eliminating confounding factors arising from landscape heterogeneity (grey bands). First row: EVI and mean annual rainfall both change along transect length. Second and third rows show spatial variance and spatial ACF-1 in EVI, respectively, along transects. We discarded regions of transects (grey bands) dominated by local heterogeneity in soil, slope, aspect or elevation (also see Fig. S13, S14, S15 in Appendix S5). In the remaining region, we identified peaks in both spatial metrics that occur within 4 km of each other as coinciding peaks (blue stars in A2, A3, B2 and B3). We estimated the associated rainfall value as the critical points (Table 1). The control Box C, which did not show bimodality, offered no estimates, consistent with the theory. Scatter data represent measurements on a moving window of 8 km *×* 8 km with a moving distance of 2 km; connecting solid line is obtained by a smoothing function with the smoothing parameter (spar) = 0.6.

**Table 1:** Estimates of the critical values of mean annual rainfall in Box-A, Box-B and Box-C from the transects using variance and ACF methods. Dash represents the transects which do not provide any estimates.

| Transect no. | Box-A |  | Box-B |  | Box-C |  |
| --- | --- | --- | --- | --- | --- | --- |
|  | CP estimated by |  | CP estimated by |  | CP estimated by |  |
|  | Variance method | ACF method | Variance method | ACF method | Variance | ACF |
| 1 | 1334 mm | 1283mm | - | - | - | - |
| 2 | - | - | 1306 mm | 1306 mm | - | - |
| 3 | 1108 mm | 1108 mm | 1281 mm | 1281 mm | - | - |

We compute the same metrics for the null transect data and show that the peaks of spatial variance and ACF-1 for Box-A and Box-B do not coincide by chance (Fig. S16 in Appendix S6). We also show that our estimations are robust to the choice of smoothing parameter (Fig. S17 in Appendix S6) and the width of the region (from 4 km to 16 km) eliminated because of landscape heterogeneity. The only exception is in Box-B for the width of heterogeneity region 12 km (or 16 km); we find that one (or both) of the critical point estimations are confounded.

#### Estimated critical points from transects in Box-A and Box-B lie close to critical points of the state diagram

The state diagrams of Box-A and Box-B show bimodality with high and low EVI values at intermediate rainfall values (Fig. 4 A,B); the occurrence of bimodality in EVI is not associated with bimodality in rainfall (Fig S6, S7 in Appendix S4).This suggests the existence of alternative stable states in these regions. Recall that critical points can be independently estimated as the threshold for the onset or disappearance of bimodality in a state diagram. For Box-A, we obtain an estimate of the critical value to be around 1000-1100 mm mm/year for the transition from high to low EVI state (Fig. 4 A); from transects, the estimated critical rainfall values range from 1108 to 1334 mm/year (Table 1). Likewise, in Box-B (Africa), the estimated critical points from the state diagram (1300-1400 mm) are close to estimates from analyses of transects (1281, 1306 mm; Table 1). We do not get any estimates from transects in the control Box-C, which is consistent with the state diagram as there is only one EVI mode at a given rainfall value (Fig. 4 C).

**Figure 4:**
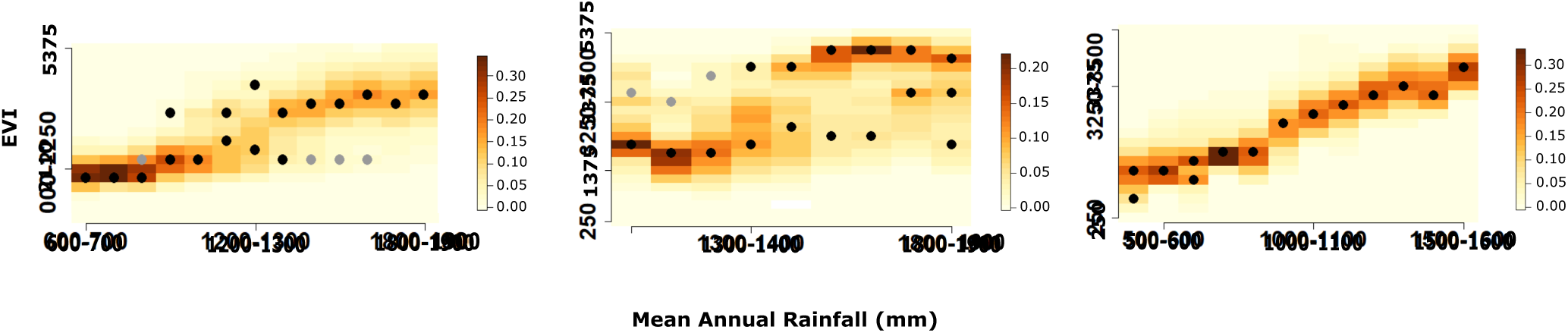
State diagrams for the three boxes. (A and B) show that Box-A and Box-B have two EVI modes occurring at comparable values of mean annual rainfall (between 1000 mm to 1300 mm in Box-A and above 1300 mm in Box-B). For each rainfall bin, rainfall does not show bimodality (see Appendix S4). These suggest evidence for alternative stable states in EVI. For each rainfall bin, black dots show the location of modes of EVI density whereas colour maps, plotted using image.plot in R, show the density of EVI. If the ratio of the density at two modes is less than 0.25, it is plotted as a grey dot. (C) does not show bimodality.

## 5 Discussion

Our analyses of spatial models showed that the ecological state and driver values corresponding to regions with simultaneous maxima of spatial variability and autocorrelations offer a quantitative estimation of critical points. We demonstrated the validity of the method using remotely-sensed vegetation data from regions in Africa and Australia. Our findings show that it is possible to estimate critical points and to identify critical regions prone to regime shifts in the future from spatial data of ecological systems.

Our method can be applied on spatial snapshots spanning alternative stable states of ecosystems even when they are in nonsteady-state conditions. A gradient of states is often maintained by an underlying gradient of a driver. Using such data, one can obtain a relation between the state of the ecosystem and the driver. Since real world data is rarely in steady state, this relationship may not show a threshold behaviour even if the underlying dynamics exhibits a critical point. We tested the applicability of our method in non-steady state conditions by simulating three scenarios: (a) a null model that exhibits no transition (Fig. 1 A), (b) a model that shows continuous transition (Fig. 1 B) and (c) a strong positive feedback model that exhibits abrupt critical transition (Fig. 1 C). Even under such circumstances masking the underlying character of the transitions, our method offers reasonable estimates of the critical points, thus showing promise for real world applications.

Our method allows estimation of critical points even with relatively small spatial datasets. We found that critical-point estimates (Table 1) from transects (8 km *×* 90 km) are comparable to those from an independent method that used data at regional scales (200 km *×* 250 km, about two orders of magnitude larger than transects). We compared our results with a previous study at a continental scale that used a different dataset (MODIS woody cover from Africa) (Staal *et al.* 2016). From Fig. 3 of Staal *et al.* (2016), we identified the critical points of transition from high to low cover to be 1300-1400 mm mean annual rainfall, comparable to our estimates. Consistency of results across scales lends credence to our claim that critical points can be quantified even with relatively small spatial datasets, offering promise of applications to ecosystems.

Is our method prone to false or failed positives? In our study, we did not find any false positives. Specifically, we chose a control region with no critical points (Box-C); none of the transects in this region provide estimations of critical points. On the other hand, our method has a failure rate. For example, one of three transects in both Box-A and Box-B (transect-2 in Box-A and transect-1 in Box-B) failed to provide any estimations of critical points. Nevertheless, even in these failed transects, both spatial variance and ACF-1 peak at the critical rainfall values expected from the state diagrams. However, those regions also occur in the vicinity of landscape heterogeneity peaks. It is difficult to disentangle whether the peak is because of the internal dynamics or the external heterogeneity and therefore, we did not consider estimates from these transects.

Ecosystems also exhibit *regular* or Turing-like patterned states, such as gaps, labyrinths or spots (Rietkerk & Van de Koppel 2008), with a characteristic length-scale of spatial variation. Such regular patterns, not considered in our study, may arise from scale-dependent processes such as short-scale positive feedback with a large-scale negative feedback (Borgogno *et al.* 2009; Meron 2012). Even in these systems, approach to critical points can be preceded by critical slowing down, simplest measures of spatial (Dakos *et al.* 2011; Kéfi *et al.* 2014). It is worth exploring, both theoretically and empirically, whether our proposed method can be applied to such pattern forming systems. This may require probing fluctuations around the characteristic scale of the pattern.

Tropical vegetation biomes show multimodality as a function of rainfall (Hirota *et al.* 2011). If we apply our method to spatial gradients that span such multiple stable states, we expect the maxima of variance and autocorrelation to occur at multiple locations, each corresponding to transition point between alternative states. Occurrence of such maxima can be confounded when the driver gradient is steep; for example, when the driver gradient exceeds a threshold value, a new type of transition, known as rate-induced tipping, can arise in dynamical systems (Ashwin *et al.* 2012; Siteur *et al.* 2016). Given that our driver gradients are modest (Fig 3 A1-C1), our inferences are possibly free of such complications.

Given the generality of the principles that underlie our method, it can be applied to a variety of ecosystems that exhibit alternative stable states. Therefore, our method enables ecosystem managers to obtain estimates of threshold or critical values of ecosystem drivers. Unlike previous qualitative early-warning indicators, our method allows for the quantitative estimation of critical points. Future research could focus on extending our methods to exploit not only spatial snapshots but also how those patterns change over time (Verbesselt *et al.* 2016; Weissmann & Shnerb 2016).

## 6 Acknowledgements

VG acknowledges support from a DBT Ramalingaswami fellowship, the DBT-IISc partnership program, and the ISRO-IISc Space Technology Cell. KT was supported by a National Postdoctoral Fellowship from SERB, Govt. of India. SR was supported in part by a J C Bose National Fellowship of the SERB, Govt. of India. Authors declare no conflicting interests.

## References

Ashwin, P., Wieczorek, S., Vitolo, R. & Cox, P. (2012). Tipping points in open systems: bifurcation, noise-induced and rate-dependent examples in the climate system. Phil. Trans. R. Soc. A, 370, 1166–1184.

Boettiger, C. & Hastings, A. (2013). Tipping points: From patterns to predictions. Nature, 493, 157–158.

Bontemps, S., Defourny, P., Bogaert, E. V., Arino, O., Kalogirou, V. & Perez, J. R. (2011). Globcover 2009-products description and validation report.

Borgogno, F., D’Odorico, P., Laio, F. & Ridolfi, L. (2009). Mathematical models of vegetation pattern formation in ecohydrology. Reviews of Geophysics, 47, RG1005.

Burthe, S. J., Henrys, P. A., Mackay, E. B., Spears, B. M., Campbell, R., Carvalho, L., Dudley, B., Gunn, I. D., Johns, D. G., Maberly, S. C. et al. (2016). Do early warning indicators consistently predict nonlinear change in long-term ecological data? Journal of Applied Ecology, 53, 666–676.

Carpenter, S. & Brock, W. (2006). Rising variance: a leading indicator of ecological transition. Ecology letters, 9, 311–318.

Carpenter, S. R., Cole, J. J., Pace, M. L., Batt, R., Brock, W., Cline, T., Coloso, J., Hodgson, J. R., Kitchell, J. F., Seekell, D. A. et al. (2011). Early warnings of regime shifts: a whole-ecosystem experiment. Science, 332, 1079–1082.

Chaikin, P. M. & Lubensky, T. C. (2000). Principles of condensed matter physics, vol. 1. Cambridge Univ Press.

Dakos, V., Kéfi, S., Rietkerk, M., Van Nes, E. H. & Scheffer, M. (2011). Slowing down in spatially patterned ecosystems at the brink of collapse. The American Naturalist, 177, E153–E166.

Dakos, V., Scheffer, M., van Nes, E. H., Brovkin, V., Petoukhov, V. & Held, H. (2008). Slowing down as an early warning signal for abrupt climate change. Proceedings of the National Academy of Sciences, 105, 14308–14312.

Dakos, V., van Nes, E. H., Donangelo, R., Fort, H. & Scheffer, M. (2010). Spatial correlation as leading indicator of catastrophic shifts. Theoretical Ecology, 3, 163–174.

Durrett, R. & Neuhauser, C. (1991). Epidemics with recovery in d= 2. The Annals of Applied Probability, 189–206.

D’Souza, K., Epureanu, B. I. & Pascual, M. (2015). Forecasting bifurcations from large perturbation recoveries in feedback ecosystems. PloS ONE, 10, e0137779.

Eby, S., Agrawal, A., Majumder, S., Dobson, A. P. & Guttal, V. (2017). Alternative stable states and spatial indicators of critical slowing down along a spatial gradient in a savanna ecosystem. Global Ecology and Biogeography, 26, DOI: 10.1111/geb.12570.

Favier, C., Aleman, J., Bremond, L., Dubois, M. A., Freycon, V. & Yangakola, J.-M. (2012). Abrupt shifts in african savanna tree cover along a climatic gradient. Global Ecology and Biogeography, 21, 787–797.

Glenn, E. P., Huete, A. R., Nagler, P. L. & Nelson, S. G. (2008). Relationship between remotely-sensed vegetation indices, canopy attributes and plant physiological processes: what vegetation indices can and cannot tell us about the landscape. Sensors, 8, 2136–2160.

Google Earth Engine Team (2015). Google earth engine: A planetary-scale geo-spatial analysis platform. URL https://earthengine.google.com.

Guttal, V. (2008). Applications of nonequilibrium statistical physics to ecological systems. Ph.D. thesis, The Ohio State University.

Guttal, V. & Jayaprakash, C. (2008). Changing skewness: an early warning signal of regime shifts in ecosystems. Ecology letters, 11, 450–460.

Guttal, V. & Jayaprakash, C. (2009). Spatial variance and spatial skewness: leading indicators of regime shifts in spatial ecological systems. Theoretical Ecology, 2, 3–12.

Guttal, V., Raghavendra, S., Goel, N. & Hoarau, Q. (2016). Lack of critical slowing down suggests that financial meltdowns are not critical transitions, yet rising variability could signal systemic risk. PloS ONE, 11, e0144198.

Hijmans, R. J., Cameron, S. E., Parra, J. L., Jones, P. G. & Jarvis, A. (2005). Very high resolution interpolated climate surfaces for global land areas. International journal of climatology, 25, 1965–1978.

Hirota, M., Holmgren, M., Van Nes, E. H. & Scheffer, M. (2011). Global resilience of tropical forest and savanna to critical transitions. Science, 334, 232–235.

Huete, A., Didan, K., Miura, T., Rodriguez, E. P., Gao, X. & Ferreira, L. G. (2002). Overview of the radiometric and biophysical performance of the modis vegetation indices. Remote sensing of environment, 83, 195–213.

Hughes, T. P. (1994). Catastrophes, phase shifts, and large-scale degradation of a caribbean coral reef. Science-AAAS-Weekly Paper Edition, 265, 1547–1551.

Kéfi, S., Guttal, V., Brock, W. A., Carpenter, S. R., Ellison, A. M., Livina, V. N., Seekell, D. A., Scheffer, M., van Nes, E. H., Dakos, V. et al. (2014). Early warning signals of ecological transitions: methods for spatial patterns. PloS ONE, 9, e92097.

Kéfi, S., Rietkerk, M., Alados, C. L., Pueyo, Y., Papanastasis, V. P., ElAich, A. & De Ruiter, P. C. (2007). Spatial vegetation patterns and imminent desertification in mediterranean arid ecosystems. Nature, 449, 213–217.

Kéfi, S., Rietkerk, M., Roy, M., Franc, A., De Ruiter, P. C. & Pascual, M. (2011). Robust scaling in ecosystems and the meltdown of patch size distributions before extinction. Ecology letters, 14, 29–35.

Lübeck, S. (2006). Tricritical directed percolation. Journal of statistical physics, 123, 193–221.

Meron, E. (2012). Pattern-formation approach to modelling spatially extended ecosystems. Ecological Modelling, 234, 70–82.

Noy-Meir, I. (1975). Stability of grazing systems: an application of predator-prey graphs. The Journal of Ecology, 63, 459–481.

Perretti, C. T. & Munch, S. B. (2012). Regime shift indicators fail under noise levels commonly observed in ecological systems. Ecological Applications, 22, 1772–1779.

Ratajczak, Z., Nippert, J. B. & Ocheltree, T. W. (2014). Abrupt transition of mesic grassland to shrubland: evidence for thresholds, alternative attractors, and regime shifts. Ecology, 95, 2633–2645.

Reed, D., Anderson, T., Dempewolf, J., Metzger, K. & Serneels, S. (2009). The spatial distribution of vegetation types in the serengeti ecosystem: the influence of rainfall and topographic relief on vegetation patch characteristics. Journal of Biogeography, 36, 770–782.

Rietkerk, M. & Van de Koppel, J. (2008). Regular pattern formation in real ecosystems. Trends in ecology & evolution, 23, 169–175.

Sankaran, M., Ratnam, J. & Hanan, N. (2008). Woody cover in african savannas: the role of resources, fire and herbivory. Global Ecology and Biogeography, 17, 236–245.

Sankaran, S., Majumder, S., Kéfi, S. & Guttal, V. (2017). Implications of being discrete and spatial for detecting early warning signals of regime shifts. Ecological Indicators.

Scheffer, M., Bascompte, J., Brock, W. A., Brovkin, V., Carpenter, S. R., Dakos, V., Held, H., Van Nes, E. H., Rietkerk, M. & Sugihara, G. (2009). Early-warning signals for critical transitions. Nature, 461, 53–59.

Scheffer, M., Carpenter, S., Foley, J. A., Folke, C. & Walker, B. (2001). Catastrophic shifts in ecosystems. Nature, 413, 591–596.

Schneider, F. D. & Kéfi, S. (2015). Spatially heterogeneous pressure raises risk of catastrophic shifts. Theoretical Ecology, 9, 207–217.

Sethna, J. (2006). Statistical mechanics: entropy, order parameters, and complexity. Oxford University Press.

Siteur, K., Eppinga, M. B., Doelman, A., Siero, E. & Rietkerk, M. (2016). Ecosystems off track: rate-induced critical transitions in ecological models. Oikos, 125, 1689–1699.

Staal, A., Dekker, S. C., Xu, C. & van Nes, E. H. (2016). Bistability, spatial interaction, and the distribution of tropical forests and savannas. Ecosystems, 19, 1080–1091.

Staver, A. C., Archibald, S. & Levin, S. A. (2011). The global extent and determinants of savanna and forest as alternative biome states. Science, 334, 230–232.

Steele, J. H. (1998). Regime shifts in marine ecosystems. Ecological Applications, 8, S33–S36.

van de Koppel, J., Rietkerk, M. & Weissing, F. J. (1997). Catastrophic vegetation shifts and soil degradation in terrestrial grazing systems. Trends in Ecology & Evolution, 12, 352–356.

Van Nes, E. H. & Scheffer, M. (2007). Slow recovery from perturbations as a generic indicator of a nearby catastrophic shift. The American Naturalist, 169, 738–747.

Verbesselt, J., Umlauf, N., Hirota, M., Holmgren, M., Van Nes, E. H., Herold, M., Zeileis, A. & Scheffer, M. (2016). Remotely sensed resilience of tropical forests. Nature Climate Change, 6, 1028–1031.

Weissmann, H. & Shnerb, N. M. (2016). Predicting catastrophic shifts. Journal of theoretical biology, 397, 128–134.

Wissel, C. (1984). A universal law of the characteristic return time near thresholds. Oecologia, 65, 101–107.

